# Think Before You List: Integrative Systematics of an ‘Endemic and Critically Endangered’ Butterfly in the Sundarbans

**DOI:** 10.1101/2025.10.07.680979

**Authors:** Md Jahir Rayhan, Sayema Jahan, Mohammad Tofazzal Hossain Howlader, Tanvir Ahmed, Sidratul Muntaaha Twaha, Md Fahim, Shafique Haider Chowdhury

**Author notes:** Corresponding author Md Jahir Rayhan McGuire Center for Lepidoptera & Biodiversity Florida Museum of Natural History University of Florida, Gainesville, FL 32611, United States.

## Abstract

Accurate species delineation is fundamental to understanding evolutionary relationships and guiding effective conservation measures. While the IUCN Red List serves as a critical conservation tool, its reliability hinges on precise systematic species identification and comprehensive distribution data. However, taxonomic inconsistencies remain a major challenge, particularly for poorly studied taxa in complex ecoregions like the Sundarbans mangrove forest. Here, we investigated the taxonomic identity of the Sundarbans Crow*, Euploea crameri nicevillei* (Moore, 1890), which is considered the only endemic and critically endangered butterfly of Bangladesh, reported only from the Sundarbans mangroves. The taxonomic status of this butterfly has long been debated, with authors regarding it as a morph of *Euploea core* (Cramer, 1780), while others designated it as a distinct subspecies of *E. crameri* (Lucas, 1853). Thus, we applied an integrative taxonomic approach incorporating morphological and molecular evidence with field observations and found that the *E. crameri nicevillei* is conspecific with the *E. core*. Based on its distinctive external wing morphology and geographic distribution, we recognize it as a valid subspecies of that, *E. core nicevillei* (Moore, 1890), **stat. nov.** Furthermore, this subspecies has a wider geographical range than previously assumed. These results suggest that its earlier designation as endemic and critically endangered was premature, emphasizing the importance of rigorous integrative systematics and the need to reassess its conservation status in Bangladesh.

## Introduction

Globally, the International Union for Conservation of Nature (IUCN) Red List serves as a powerful and comprehensive resource for the conservation of threatened organisms (Rodrigues 2006). However, effective conservation depends on the accurate recognition of conservation units, such as species, highlighting the critical role of proper systematics (Mace 2004; Singh 2025). Without a robust taxonomic framework, conservation efforts are likely to fail, potentially causing devastating and irreversible impacts on global biodiversity (Thomson et al. 2018). Despite this, taxonomic inconsistencies remain a significant challenge, particularly in lineages with subtle morphological differences, cryptic diversity, unresolved phylogenies, or those inhabiting poorly studied and complex ecoregions (Singh 2025).

Bangladesh lies at the crossroads of the Indo-Himalayan and Indo-Chinese biodiversity realms, making its biodiversity exceptionally rich (IUCN Bangladesh 2014). Among the insect fauna, butterflies are relatively well documented compared to other invertebrates in the country. In 2015, the IUCN Bangladesh Red List assessed a total of 305 butterfly species, of which 188 were categorized as threatened, out of at least 421 species reported to date (Hossain 2023). The only taxon listed as Critically Endangered is a distinctive nymphalid subspecies, the Sundarbans Crow, *Euploea crameri nicevillei* (Moore, 1890). This taxon has also been claimed to be the only endemic butterfly of Bangladesh, occurring exclusively in the Bangladesh portion of the Sundarbans mangrove forest (Chowdhury 2015; Hossain 2013). Earlier reports suggested that this subspecies had a wide distribution throughout the forest, but more recent accounts indicate that its range has contracted to an area of only 78.54 km², allegedly due to a variety of largely unknown factors (Chowdhury 2015).

Historically, *Euploea crameri nicevillei* has been considered rare in multiple accounts (Moore 1890; Talbot 1947). The original voucher specimens were identified as the morphotype of *Euploea vermiculata* (Moore 1890), and then it was designated as a distinct taxon– *Tronga nicevillei* (Moore 1890). Meanwhile, it was also suggested to be a dry season form of the more common species *Euploea core* (Cramer, 1780). But, both *Euploea core* and *Euploea crameri nicevillei* morphs are sympatric in the Sundarbans mangrove forest (Larsen 2004).

The tangled history of this butterfly motivated our investigation, driven by two central questions: what exactly is *Euploea crameri nicevillei* (Moore, 1890) and does it truly merit the recognition as an endemic critically endangered taxon in Bangladesh? We conducted a comprehensive investigation of reviewing historical literatures, type specimens and in-field samplings, with morphological and molecular characterization to address these questions.

## Materials and methods

### Sample Collection

Under the permission of the Bangladesh Forest Department (source letter number: 22.01.0000.004.04.21.1.23.948), we conducted three consecutive field expeditions between November 2024 and August 2025 in the Sundarbans Mangrove Forest of Bangladesh. Our sampling sites included Katka and Kachikhali (Sarankhola administrative range) and Harbaria and Baiddomari (Chandpai administrative range). At each study site, five observers conducted surveys of the Sundarbans Crow butterfly along 500 m transects established on pre-marked walking trails. All flowering mangrove plants along these transects were examined to record butterfly presence and activity. Additionally, mangrove-dwelling Apocynaceae plants were inspected for potential larval occurrences. Observations included the behavioral and ecological traits of the target species to support assessments of its distribution and host-plant associations. When the target species was encountered, we recorded the number of individuals and the GPS location for each observation. Using sweep nets, we captured a total of five Sundarbans Crow and five Common Crow specimens from the study sites. We then carefully removed legs from Sundarbans Crow specimens (n=5) using sterilized forceps and gloves to prevent contamination and stored them in 95% ethanol with proper labeling. The remaining specimens (n=5) were euthanized using acetone vapor, then pinned and spread following standard entomological procedures for morphological examination. The specimens are deposited at the Invertebrate Museum of the Department of Zoology, University of Chittagong.

### Morphological Analysis

Genitalia dissections were performed with minor modifications following Robinson (1976). After detachment of the genital capsules from the abdomen boiled in 10% KOH (for how long?), they were treated with sequential grades of alcohol from 30–100%, and later were kept submerged in clove oil for 2–3 hours to get the soft tissues digested. Permanent slides were mounted on DPX medium and cover-slipped. Genital structures were examined and photographed under an L 101 compound microscope, while adult specimens were imaged using a Nikon D5500 DSLR camera equipped with an 18–55 mm lens. In addition, type specimens of *Tronga nicevillei* Moore, 1890 deposited in the Natural History Museum in London, UK (NHM, UK), were examined to ensure precise comparison.

### Molecular Analysis

DNA extraction, PCR, Sequencing: Genomic DNA was extracted from leg samples preserved in ethanol using the DNeasy Blood and Tissue Kit (Qiagen, Germany) following the manufacturer’s protocol. Briefly, leg samples were crushed in ATL buffer, incubate for lysis and the total genomic DNA was extracted using DNeasy Mini spin column. The concentration of eluted DNA was measured using nanodrop spectrophotometer. We used two individuals for the purpose (labels: H250826-025 I06 JS1_Tronga_nicevillei, and H250826-025 M06 JS3_Tronga_nicevillei). The mitochondrial Cytochrome C Oxidase subunit I (COI) gene was amplified using the primer pair LepF1 and LepR1 (Hebert at al. 2004) in a 25 μL reaction volume carried out in a thermal cycler (SimpliAmp^TM^Thermal Cycler, Applied Biosystem). The PCR protocol consisted of an initial denaturation at 94 °C for 5 min, followed by 40 cycles of denaturation at 94 °C for 30 s, annealing at 50 °C for 30 s, and extension at 72 °C for 90 s, with a final extension at 72 °C for 10 min. The amplification yielded a ∼650 bp fragment of the COI gene, which was subsequently purified and sent to Macrogen Inc. (Seoul, South Korea) for bidirectional Sanger sequencing. The quality of raw sequences was assessed using BioEdit v7.2 (Hall, 1999). Sequence alignment, editing, and generation of consensus sequences were performed in Geneious v11.1.5 (Kearse et al., 2012). The sequences have been deposited in GenBank under accession numbers PX431589 and PX431590 (currently listed under *Euploea crameri nicevillei*; corresponding to *Euploea core nicevillei*, **stat. nov.** as treated herein).

### Phylogenetic analysis

Phylogenetic relationships among the obtained COI sequences were inferred using both Neighbor-Joining (NJ) and Maximum Likelihood (ML) approaches. Additional sequences for other *Euploea* species were retrieved from NCBI GenBank following a BLAST search (Accession numbers: MF804643.1, PP405031.1, LC819331.1, GU012616.1, ON075551.1, KR150520.1, PP726767.1, PQ524941.1, KT879888.1, KR150622.1). Additionally, three COI sequences previously deposited in GenBank by Hossain *et al*. (unpublished, direct submission) under the name “*Euploea crameri nicevillei*” (Accession numbers: PV409685.1, MH269417.1, PV592280.1) from Bangladesh were included in our analyses. The cytochrome C oxidase I (COI) gene sequence of *Danaus chrysippus* (Accession number: LC877962.1) was used as the outgroup.

Pairwise sequence alignment was performed using the MUSCLE algorithm in MEGA X (Kumar et al., 2018), and genetic distances were calculated under the Kimura 2-parameter (K2P) model (Kimura, 1980). The alignments were checked visually, and both ends of the sequence were trimmed to avoid low-quality base pairs. To assess the potential impact of multiple substitutions on phylogenetic inference, the aligned COI sequences were tested for substitution saturation using the Xia et al. (2003) method implemented in DAMBE 6. The test calculates the index of substitution saturation (Iss) and compares it to a critical value (Iss.c) for both symmetrical and asymmetrical tree topologies. Sequences with Iss significantly lower than Iss.c are considered suitable for phylogenetic analysis, indicating that the alignment retains sufficient phylogenetic signal despite potential multiple substitutions. The NJ tree was reconstructed in MEGA X using K2P distances, with branch support assessed by 1,000 bootstrap replicates. Besides, the ML tree was inferred using IQ-TREE v2 on the aligned nucleotide sequences.

ModelFinder (-m MFP) selected the best-fit substitution model: GTR+F+G4 according to the AIC and corrected AIC, and TIM2+F+I according to BIC, with the latter used for tree reconstruction. Branch support was evaluated using 1,000 replicates each of the ultrafast bootstrap approximation (UFBoot) (-B 1000) and the Shimodaira–Hasegawa-like approximate likelihood ratio test (SH-aLRT) (-alrt 1000). UFBoot provides fast, accurate bootstrap support (Hoang et al., 2018), while SH-aLRT assesses branch reliability by comparing the likelihood of alternative topologies (Guindon et al. 2010). Analyses utilized all available CPU cores (–T AUTO), and resulting trees were visualized and annotated in FigTree v1.4.4 (Rambaut 2018), and further edited for publication-quality presentation in Inkscape.

## Results

### Historic overview of Euploea species

The famous lepidopterist Charles Lionel Augustus de Nicéville obtained four specimens of a danaid butterfly from the Sundarbans near Kolkata, India, although the exact date of collection was not recorded. de Nicéville (1901) only remarked that “many years ago four specimens of the species were given to me, taken in February.” He initially recognized the morphotype as *Crastia vermiculata* and labeled it accordingly (Fig. 1C-D).

**Figure 1.**
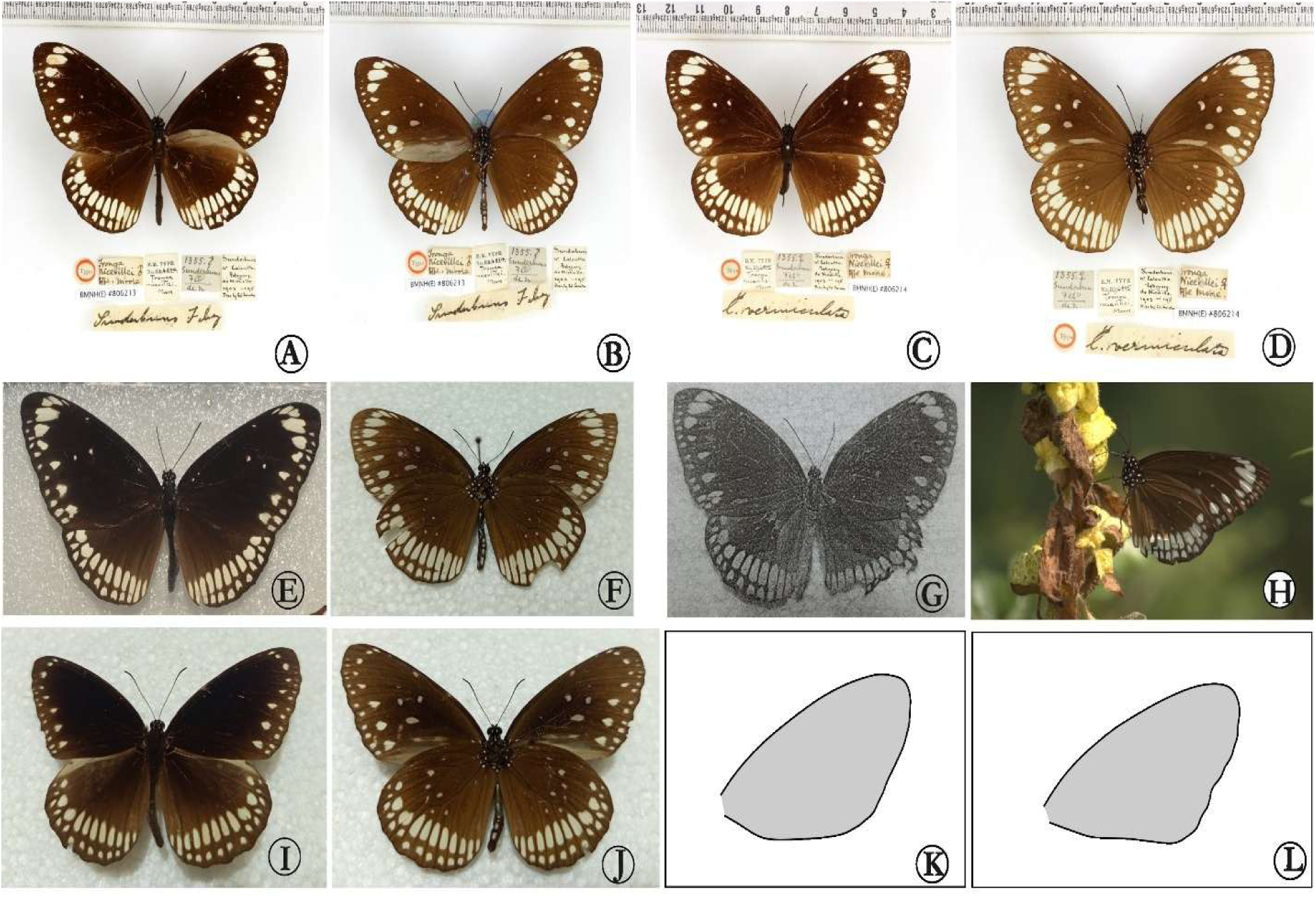
The *Euploea species*; **A.** Male type of *Tronga nicevillei* Moore, 1890, dorsal from NHM, UK; **B.** Ditto, ventral; **C.** Female type of *Tronga nicevillei* Moore, 1890 from NHM, UK; **D.** Ditto, ventral; **E.** Male *Euploea core nicevillei* (Moore, 1890), **stat. nov.** from Baiddomari, Sundarbans, Bangladesh, dorsal; **F.** Ditto, ventral; **G.** Male *Euploea core nicevillei* (Moore, 1890) from Chowdhury, 2004; **H.** *Euploea core nicevillei* (Moore, 1890) pheromone replenishing on *Crotalaria* plants for pollens from Kachikhali, Sundarbans, Bangladesh [Photo credit: Ripon Chandra]; **I.** Male *Euploea core core* (Cramer, 1780) from Katka, Sundarbans, Bangladesh, dorsal; **J.** Ditto, ventral; **K.** Male forewing shape of *Euploea core nicevillei* (Moore, 1890), **stat. nov.**, **L.** Male forewing shape of *Euploea core core* (Cramer, 1780).

Of these, two specimens were later given to Colonel Swinhoe, based on which Moore (1890) described the morphotype as *Tronga nicevillei* (Fig. 1C-D). We were unable to locate the rest two specimens and their depository. The form is unusual in three respects: (1) males lack the sexual brand in the submedian interspace of the forewing upperside; (2) the wing spots are unusually large and distinct compared to other Indian *Euploea* species; and (3) the wings are somewhat broader in shape.

Despite these discrepancies, de Nicéville (1901) maintained his original view when reviewing Indian butterflies of the *Tronga* group and treated *Limnas mutabilis cora* Hübner, 1806, *Euploea vermiculata* Butler, 1866, and *Tronga nicevillei* Moore, 1890 as three forms of *Euploea core* (Cramer, 1780). He concluded that *T. nicevillei* represented unusually white dry-season forms of *E. core* from the Sundarbans. Although acknowledging differences such as broader wings and the absence of the male brand, he doubted that the taxon was a distinct species, arguing that the Sundarbans, being a relatively recent alluvial formation, was unlikely to have produced an entirely separate species. Instead, he regarded the specimens as local variants of *E. core*.

Fruhstorfer (1904) disagreed, affirming that Moore’s (1890) treatment was correct. Later, Fruhstorfer (1910) considered *Tronga nicevillei* Moore, 1890 to be a subspecies of *Euploea crameri* Lucas, 1853, commenting (translated from German): “*nicevillei* Moore, originally described from the Sunderbunds [Sundarbans], small alluvial islands at the mouth of the Ganges, was most likely introduced there accidentally by ship traffic. It is characterized by unusually broad, pure white submarginal spots on all wings. To date, only a single female specimen is known, housed in the British Museum.”

Fruhstorfer’s claim is apparently incorrect in two key respects. First, the Sundarbans form a continuous extension of the Bengal mainland; thus, accidental introduction and speciation through isolation are improbable. Second, Moore (1890) clearly reported two specimens in the British Museum (one male and one female), which we also examined. It appears that Fruhstorfer overlooked the male specimen and based his conclusion solely on the female. Subsequently, Hulstaert (1931) and Evans (1932) followed Fruhstorfer (1910) in treating *T. nicevillei* as a subspecies of *E. crameri*. Talbot (1947) reinforced this view. Notably, all these authors examined only the two type specimens in the British Museum; none observed or obtained additional specimens, and each reported the taxon as “very rare.” After more than a century, Chowdhury (2004) rediscovered the butterfly in the Kachikhali and Jamtala regions of the Bangladesh counterparts of Sundarbans. However, Larsen (2004) noted that he had already found the species to be “quite common” in December 2002 on the Khotka Plains (Katka, Sundarbans), where it occurred alongside *E. core*. While Chowdhury (2004) described this as a rediscovery after nearly 119 years, Larsen (2004) remarked that it had also been recorded from the Orissa mangroves about a century earlier. Interestingly, IUCN (2015) listed this species as the only butterfly from Bangladesh categorized as both “Endemic” and “Critically Endangered,” based on two main considerations: (1) preliminary records at the time indicated an area of occupancy of 78.54 km², and (2) there were no recent records from the Indian part of the Sundarbans. Nevertheless, Chowdhury (2014) reported *Euploea crameri* [apparently the *Tronga nicevillei* form] from the Sundarban Biosphere Reserve, West Bengal, India, and noted its occurrence in both mangrove forests and reclaimed non-forest areas, with its status recorded as very rare.

### A distinct morph of Euploea *core* or a subspecies of Euploea crameri nicevillei?

#### Morphological evidence

As mentioned earlier, the butterfly from the Sundarbans mangroves is unusual in its external morphology, particularly in possessing broader white wing markings (Fig. 1A-H, K). The confusing, unique and distinct wing patterns and overall appearance have long puzzled lepidopterists, as discussed previously.

Comparison with *E. crameri*: *E. crameri* is a highly variable species in terms of external wing pattern, a variation that corresponds to (though not limited to) geographic distribution. The species is divided into over fifteen subspecies across the Indo-Malay region. Despite this variability, all subspecies share several consistent diagnostic features. These include: in males, the underside of the forewing bears two moderately long and broad blackish sex stripes in space 1b, arranged to form a parallelogram, with the anterior distal edge positioned closer to the termen. Additionally, the posterior half of space 1b and the whole of space 1a are greyish-white (Corbet 1942) (Fig. 2A-B). On the upper side of the forewings, males lack any distinct androconial “sex brand” (Talbot 1947).

**Figure 2.**
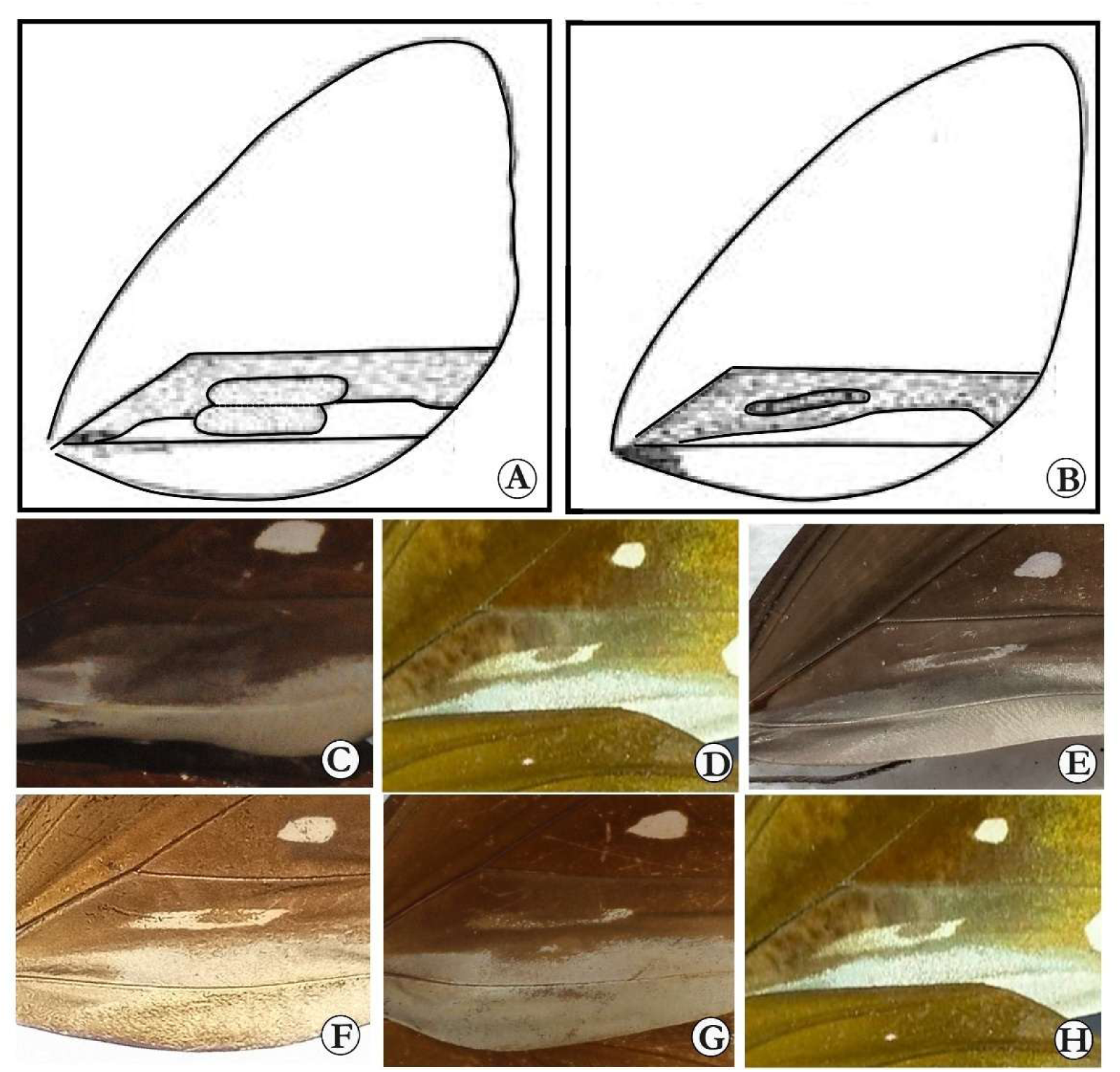
Comparisons of sex stripes on underside forewings in male. A. *Euploea crameri* Lucas, 1853 (after Corbet, 1942); B. *Euploea core* (Cramer, 1870) (ditto); C. *Euploea crameri* (from: https://commons.wikimedia.org/wiki/File:EuploeaCrameriMUpUn.jpg); D. *Euploea core core*; E. *Euploea core core*; F. *Euploea core nicevillei*, **stat. nov.**; G. *Tronga nicevillei* male type from NHM, UK; H. *Euploea core nicevillei*, **stat. nov.**

Although *Tronga nicevillei* similarly lacks the androconial “sex brand” on the upperside of male forewings, its underside sex stripes are never comparable to those of *E. crameri* (Fig. 2C-D). It is therefore evident that the historical recognition of *T. nicevillei* as a subspecies of *E. crameri* was based largely on the absence of the androconial “sex brand.” However, detailed examination reveals that the underside sex stripes of *T. nicevillei*, while superficially confusing, are more similar to those of *E. core* rather than *E. crameri*. Moreover, the absence of the “sex brand” in *E. crameri* cannot be considered a reliable diagnostic feature, as in rare cases individuals of *E. crameri* may develop such markings (Inayoshi 2025). Besides, the male genitalia of the *Tronga nicevillei* morphotype are identical to those of *E. core* rather than *E. crameri* in all respects, particularly in the shape of the valva and the cornutus (Fig. 3A-G, 4A-B).

**Figure 3.**
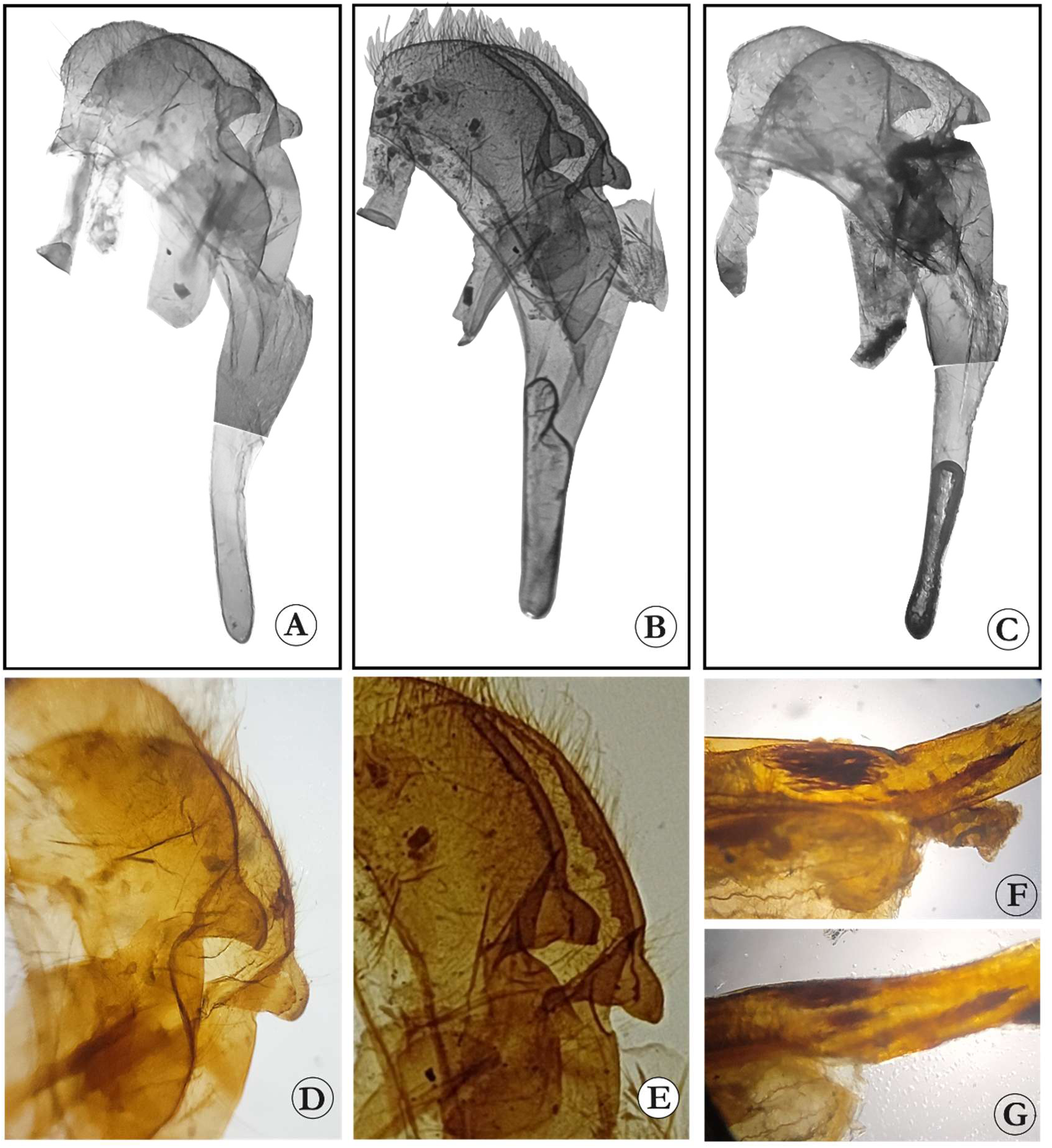
Male genitalia of *Euploea* species. A. *Euploea core core* (Cramer, 1780); B. *Euploea core nicevillei* (Moore, 1890), **stat. nov.** (JS1); C. Ditto (JS3); D. *Euploea core core* (Cramer, 1780) valva in detail; E. *Euploea core nicevillei* (Moore, 1890), **stat. nov.** (JS1) valva in detail; F. *Euploea core core* (Cramer, 1780) aedeagus in detail; G. *Euploea core nicevillei* (Moore, 1890), **stat. nov.** (JS1) aedeagus in detail.

**Figure 4.**
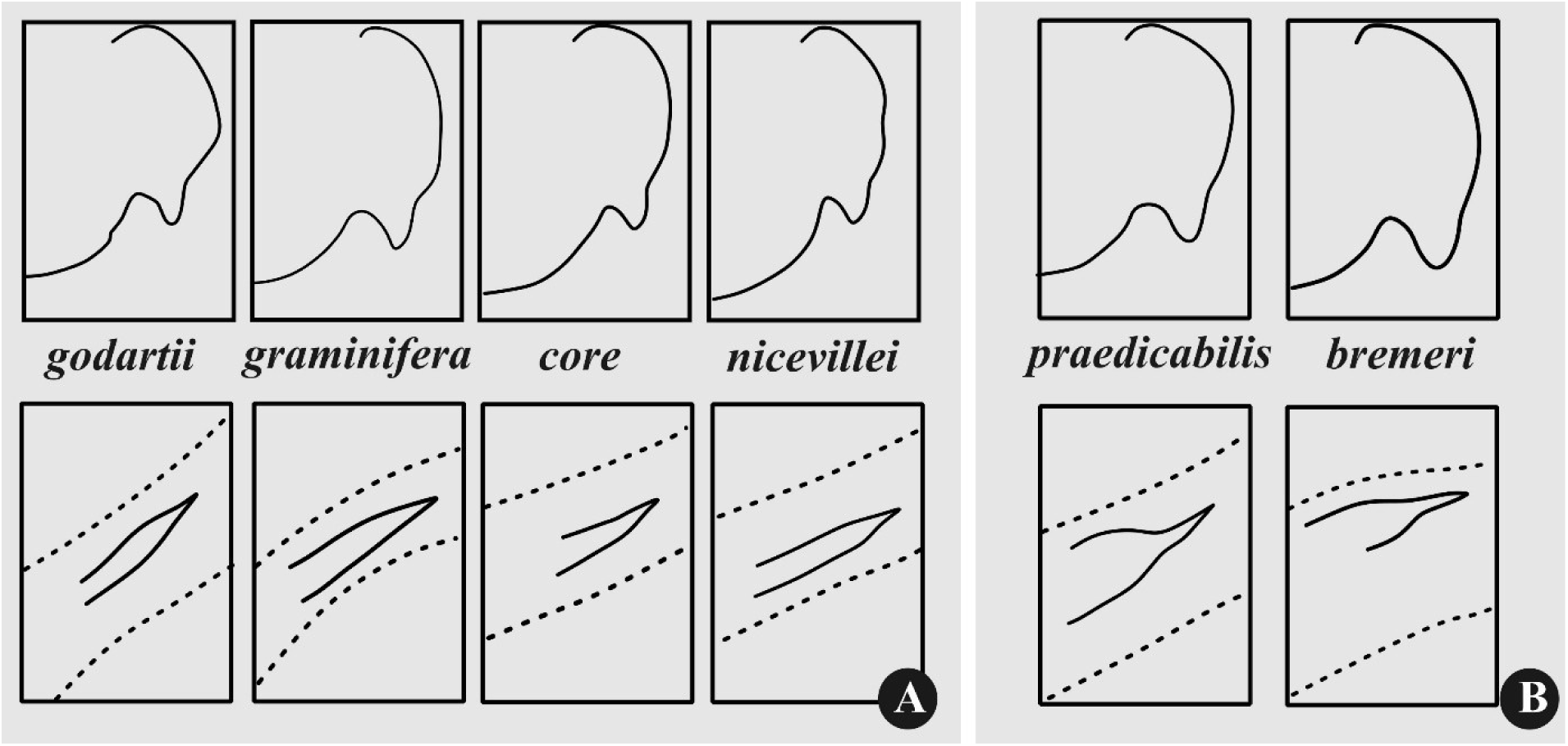
Comparisons of male genitalia. A. Different subspecies of *Euploea core*; B. Different subspecies of *Euploea crameri*. In each panel, upper one represents the right valva in detail, and the lower one represents the aedeagus in detail showing the cornutus (except *core* and *nicevillei*, rest are after Inayoshi 2025).

Comparison with *E. core*: de Niceville (1901) was a strong proponent of placing the *Tronga nicevillei* morphotype within *E. core*. The latter is characterized in males by the upperside forewing having a single sex brand, and the underside forewing displaying a pale, elongate, narrow stripe in the anterior half of space 1b. This stripe is typically overlaid by a blackish stripe, which usually obscures much of the pale marking. The posterior half of space 1b and the entirety of space 1a are shining greyish white (Corbet 1942). However, wing spots are variable, and, in some individuals, the spots are very large closely resembling those of the mangrove *Tronga nicevillei* form.

Although the *T. nicevillei* morphotype lacks the sex brand on the upperside forewings of males, its underside markings are indistinguishable from those of *E. core* (Fig. 2D-H). The upper side markings, though considered a defining feature of this species, can be obscure or even completely absent in some morphotypes (de Niceville 1901; Inayoshi 2025). This suggests that upperside characters are unreliable, whereas underside markings remain more consistent. In male genitalia, as discussed earlier, *T. nicevillei* is indistinguishable in all respects from this species.

#### Molecular Evidence

Genetic distance: The molecular data available for most *Euploea* species are limited to the universal DNA barcode locus, the mitochondrial cytochrome c oxidase subunit I (COI) gene. Accordingly, our analysis was restricted to this marker. A comparative assessment of COI sequences among *Euploea* species revealed that the evolutionary (genetic) distance between *E. core* and *Tronga nicevillei*, as estimated under the Kimura 2-parameter (K2P) model, is 0-0.2% (Table 1). This negligible divergence does not support the recognition of *Tronga nicevillei* as distinct from *E. core*.

Phylogenetic reconstruction: The Xia test of substitution saturation, performed in DAMBE (Xia et al. 2003; Xia and Lemey 2009), yielded an observed index of substitution saturation (Iss = 0.0754), which is significantly lower than the critical values for both symmetrical (Iss.c = 0.7379) and extreme asymmetrical trees (Iss.c = 0.6117; p < 0.001), indicating little saturation. This suggests that the sequences retain sufficient phylogenetic signal and are suitable for phylogenetic analysis.

As mentioned earlier, we reconstructed both Neighbor-Joining (NJ) and Maximum Likelihood (ML) trees. Although the relationships among species within the genus *Euploea* differed between analyses (likely due to the use of a single marker) (Fig 5, 6), both approaches consistently placed *Tronga nicevillei* within the *E. core* clade, with UFBoot support of 93.3 and SH-aLRT of 97 (Fig. 5). Besides, the previously generated COI sequences by Hossain *et al*. submitted to GenBank as ‘*Euploea crameri nicevillei*’ were also found to be identical to both *E. core* and the *Tronga nicevillei* morphotypes sequenced in this study, clustering within the *E. core* clade. These results further suggest that *Tronga nicevillei*, or *Euploea crameri nicevillei*, is conspecific with *E. core*.

**Figure 5.**
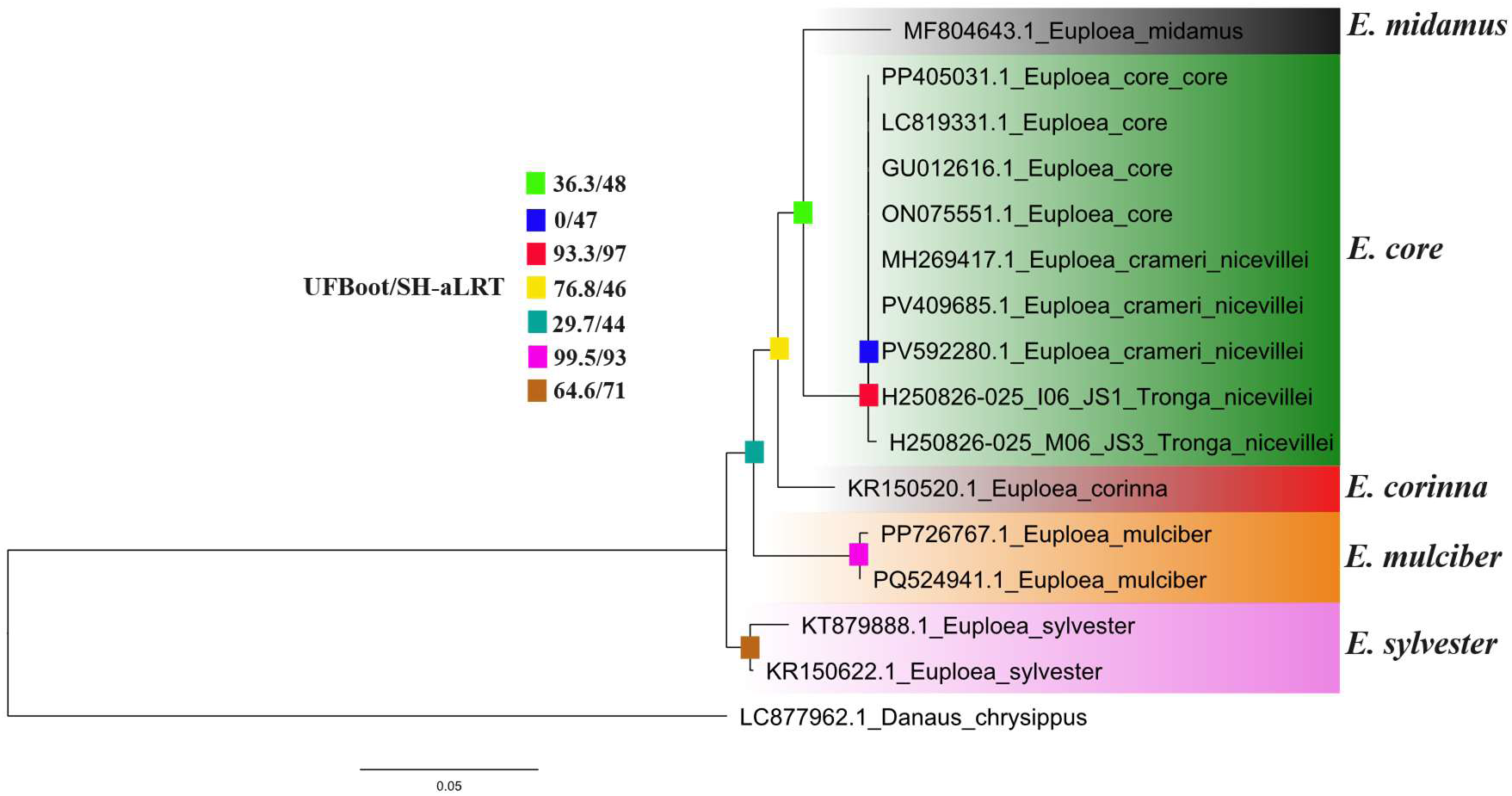
Maximum likelihood (ML) phylogeny of *Euploea* species reconstructed in IQ-TREE using the mitochondrial cytochrome oxidase I (COI) gene. Node support is shown as ultrafast bootstrap (UFBoot) values and Shimodaira–Hasegawa approximate likelihood ratio test (SH-aLRT) values. GenBank accession numbers are provided along with the scientific names.

**Figure 6.**
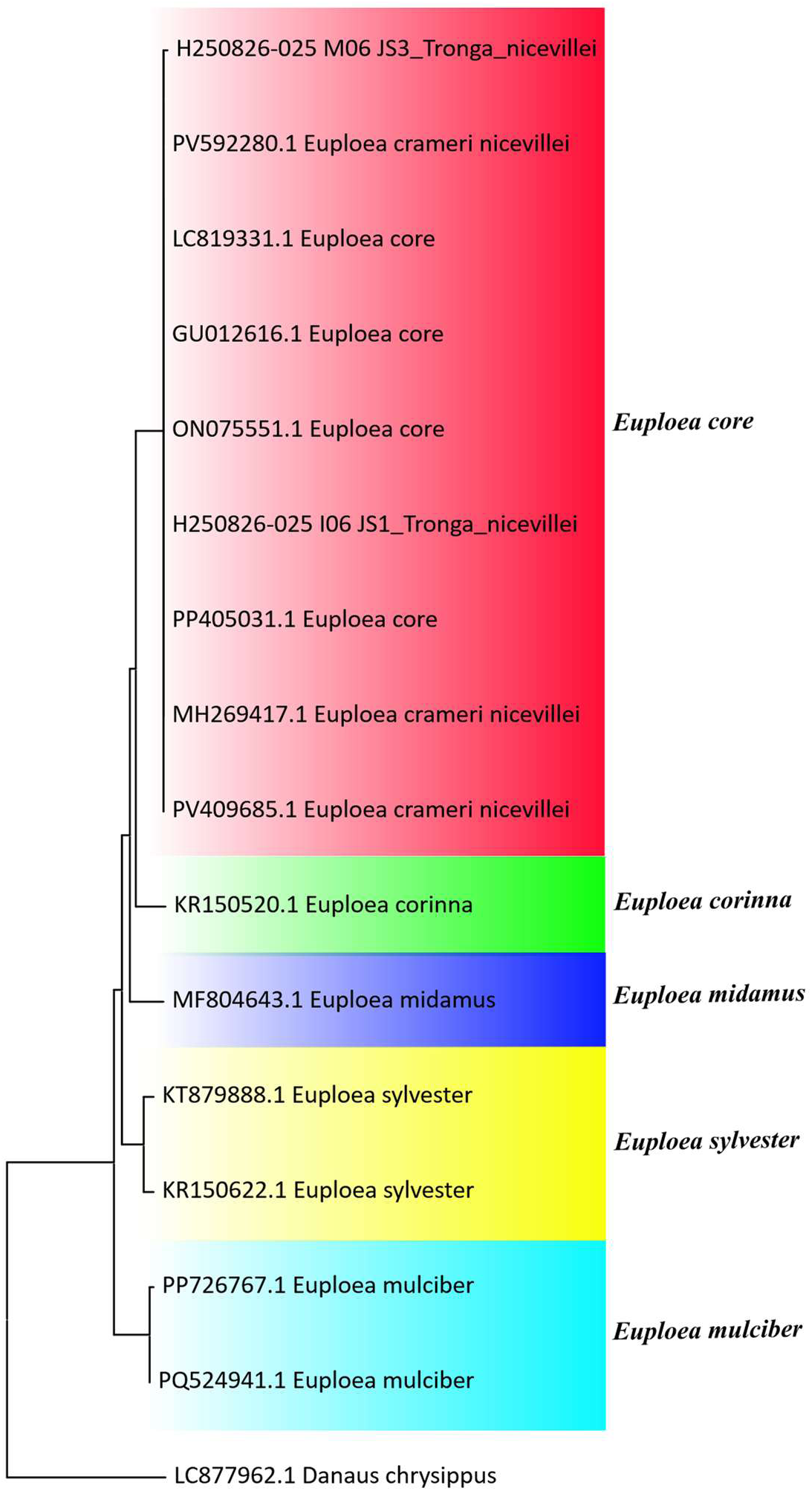
Neighbor-joining (NJ) phylogeny of *Euploea* species based on the mitochondrial cytochrome oxidase I (COI) gene, reconstructed in MEGA X. All nodes are fully supported (100%) in bootstrap analysis. GenBank accession numbers are provided along with the scientific names.

### New Systematic Account

*Euploea core nicevillei* (Moore, 1890), **stat. nov.**

*Tronga nicevillei* Moore, [1890]: 77

*=Crastia vermiculata;* Moore, [1890] (*sensu* de Nicéville)

= *Euploea crameri nicevillei* (Moore, 1890) (*sensu* Fruhstorfer, 1910; Hulstaert, 1931; Evans, 1932; Talbot, 1947)

Diagnosis: This subspecies differs from the nominotypical *E. core core* in having broader wings with larger and more distinctive markings, and in the absence of the male sex brand on the forewings upperside. These traits are consistent and readily diagnosable.

Remarks: Although the *Tronga nicevillei* form exhibits several distinguishing traits in external morphology, it shares fundamental characteristics with *Euploea core*, including the underside forewing features and male external genitalia. While the external male genitalia of *Euploea* species are generally highly conserved, subtle genital differences can still be observed among different species (Corbet 1942; Lambkin 2017). However, the genital morphology of ‘*Euploea crameri nicevillei*’ and *Euploea core* were identical with no significant difference, further supporting the conclusion that these morphs are not reproductively isolated. This supports its close affinity and likely conspecific status with *E. core*, rather than with *E. crameri*. Molecular analysis of the COI gene further corroborates this relationship.

In Katka and Kachikhali, we observed the *T. nicevillei* form occurring in sympatry with the nominal *E. core* form (Fig. 7), consistent with the earlier observations of Larsen (2004). Beyond this area, however, *E. core* was neither detected in our surveys nor reported in previous studies, suggesting that *Euploea core nicevillei*, **stat. nov.**, represents a distinct subspecies, geographically confined to the Mangroves of Bengal delta (and probably Orissa, India). Furthermore, the *T. nicevillei* form was recorded during both the dry and wet seasons, indicating that the form is stable and not a seasonal variant as indicated by de Nicéville (1901).

**Figure 7.**
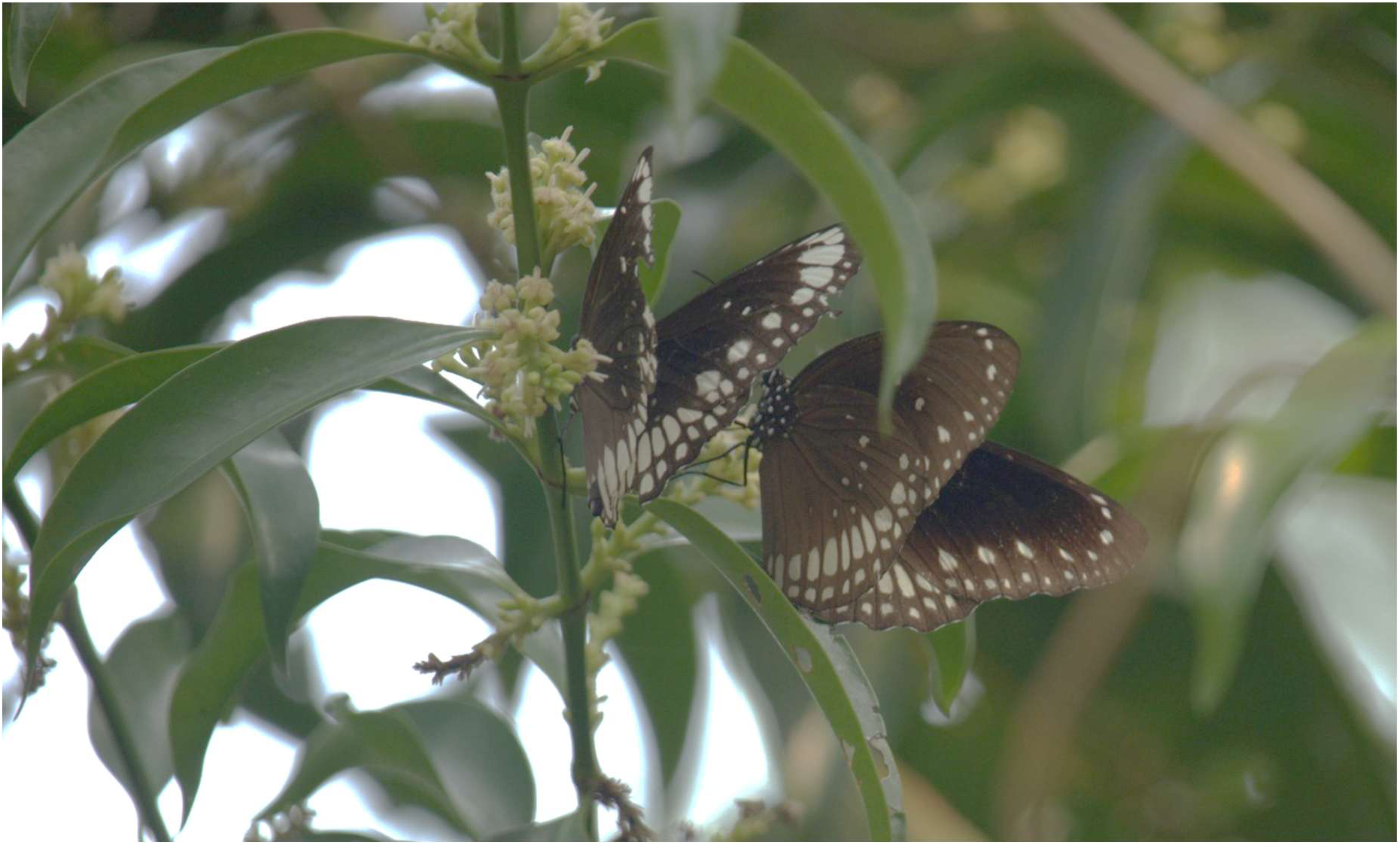
*Euploea core nicevillei*, **stat. nov.** (left) and *Euploea core core* nectaring together on *Lumnitzera racemosa* plants in Katka, Sundarbans

The distinctiveness in wing shape and pattern of the *T. nicevillei* form, together with its restricted distribution in the Sundarbans, supports recognition of this taxon as a subspecies of *E. core* rather than of *E. crameri*. The known distribution of other subspecies of *E. crameri* does not justify treating *T. nicevillei* under that species, as such an arrangement obscures the dispersal history of the Sundarbans population.

Although Fruhstorfer (1910) suggested a hypothesis involving introduction through ship trafficking, this lacks supporting evidence and cannot be considered plausible. We therefore designate this subspecies as *Euploea core nicevillei* (Moore, 1890), **stat. nov**. Katka and Kachikhali appear to represent a transitional zone at the boundary between the Sundarbans mangrove forest and the adjacent mainland, where both *E. core core* and *E. core nicevillei* **stat. nov.** occur.

Furthermore, although male genitalia and COI sequence analysis do not show significant divergence from the nominotypical form, this is not uncommon in recently diverged populations (Harris *et al*. 2018). The consistent morphological distinctiveness and restricted geographic distribution, despite limited genetic differentiation, supports treatment of this taxon as a valid subspecies rather than simply synonymizing *T. nicevillei* with *E. core*.

*Conservation Status of Euploea core nicevillei (Moore, 1890), **stat. nov***.

IUCN Bangladesh (2015) recognized *Euploea core nicevillei*, **stat. nov.** as a “Critically Endangered” and “Endemic” species in Bangladesh, which was based on CR B1ab(ii) Red List Category and Criteria (ver. 3.1; Chowdhury 2015). This means that the butterfly has a very limited range, occurs in one or very few locations, and its Area of Occupancy (AOO) is declining. Practically, the reported Extent of Occurrence (EOO) and Area of Occupancy of 79 km² is likely stemmed from extremely limited field expeditions. Although this morphologically distinct butterfly taxon has reportedly never been observed outside the mangroves of the Sundarbans and possibly Odisha in India (Larsen 2004), our recent surveys, including records from Chandpai Range, Harbaria, and Baiddomari, as well as photographic evidence from adjacent non-mangrove areas (https://www.inaturalist.org/observations/255150551), indicate that the subspecies’ range of occupancy is considerably broader than previously documented. Furthermore, reports from the Indian Sundarbans (Chowdhury 2014) and potentially from the coastal mangroves of Orissa in India may also indicate that the butterfly is distributed beyond the Bangladesh Sundarbans and can’t be endemic to Bangladesh (Fig. 8).

**Figure 8.**
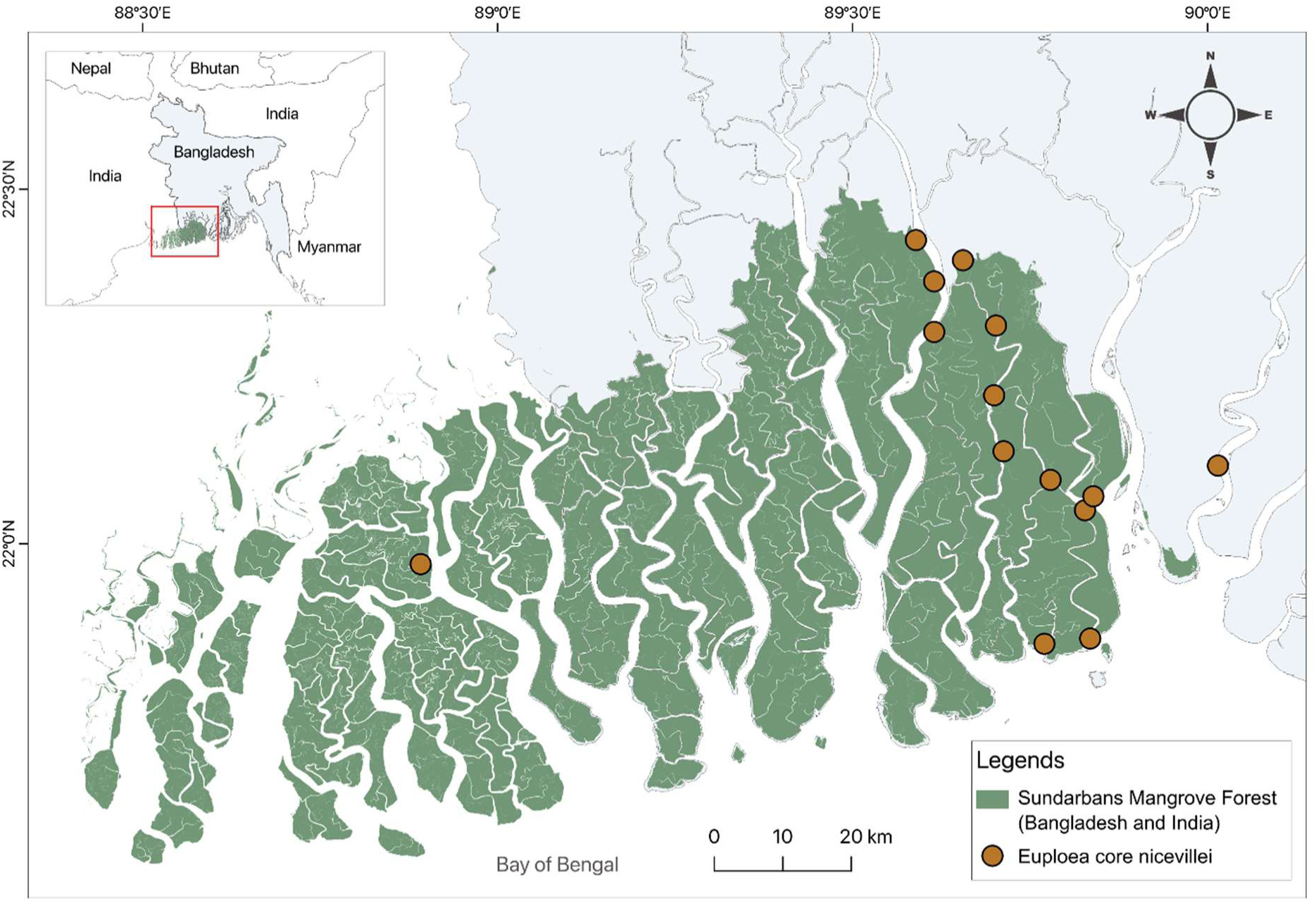
Map of Sundarbans mangrove showing the confirmed occurrence record of *E. core nicevillei* (Moore, 1890), **stat. nov.**

The Sundarbans present a challenging environment for fieldwork due to logistical difficulties and the physical demands of surveying. Consequently, butterfly studies in this region remain limited with only a few expert efforts to date, resulting in only 112 species being formally known from the Bangladesh Sundarbans (Hossain 2014; Ahmed and Mondal 2016; BFD et al. 2018), compared to 137 species in the Indian portion (Das et al. 2019). Comprehensive field surveys are therefore required for a proper assessment of the butterfly diversity in the Sundarbans.

Remarkably, we observed *E. core nicevillei* to be locally common, particularly in Katka and Baiddomari. Although the species may be rare in some areas, there is currently no evidence of rapid population decline, and further investigation is needed. Nevertheless, the Sundarbans are under severe threat from various anthropogenic pressures and climate change. Given its unique adaptation to mangrove habitats, *E. core nicevillei* merits particular conservation attention and the designation of an appropriate protected status. We strongly encourage comprehensive studies on the butterfly fauna in the Sundarbans with particular emphasis on this taxon to reassess its conservation status and develop effective conservation strategies.

## Conclusion

Our study resolves the long-standing taxonomic ambiguity of the Sundarbans Crow, confirming it as a valid subspecies of *Euploea core* rather than *E. crameri*. This study underscores the importance of integrative systematics for accurate species recognition, clarifying species boundaries and highlights the need to reassess the conservation status of this unique mangrove butterfly in Bangladesh.

## Acknowledgements

We gratefully acknowledge the Ministry of Forest and Environment, Bangladesh, for granting the necessary permission to conduct this research (Reference:22.01.0000.004.04.21.1.23.948; Date: 12 December 2024). We are also thankful to the local forest officials for their valuable support. Our sincere appreciation goes to Sumaiya Hafiz, Ripon Chandra, and Ritu Roy for their assistance during the initial field visit. We also extend our gratitude to Mr. Peter Smetacek for his insightful suggestions on systematics. TA acknowledges support from the German Academic Exchange Service (DAAD) through a doctoral research scholarship (no. 57645448). MJR and SJ especially acknowledge the help and support of Dr. Mohammad Abdul Wahed Chowdhury and Dr. Sabir Bin Muzaffar for their valuable assistance in various aspects of the project. Finally, we are deeply grateful to Mr. Mark J. Sterling from NHM, UK, for kindly sharing the photographs of type specimens.

---

**Table:** Estimates of Evolutionary Divergence between Sequences based on the Kimura 2-parameter model.

|  | 1. | 2. | 3. | 4. | 5. | 6. | 7. | 8. | 9. | 10. | 11. | 12. | 13. | 14. | 15. | 16. |
| --- | --- | --- | --- | --- | --- | --- | --- | --- | --- | --- | --- | --- | --- | --- | --- | --- |
| 1. LC877962.1_Danaus_chrysippus |  |  |  |  |  |  |  |  |  |  |  |  |  |  |  |  |
| 2. MF804643.1_Euploea midamus | 0.141 |  |  |  |  |  |  |  |  |  |  |  |  |  |  |  |
| 3. PP726767.1_Euploea mulciber | 0.149 | 0.044 |  |  |  |  |  |  |  |  |  |  |  |  |  |  |
| 4. PP405031.1_Euploea core | 0.153 | 0.032 | 0.042 |  |  |  |  |  |  |  |  |  |  |  |  |  |
| 5. PQ524941.1_Euploea mulciber | 0.146 | 0.046 | 0.002 | 0.040 |  |  |  |  |  |  |  |  |  |  |  |  |
| 6. KR150520.1_Euploea corinna | 0.161 | 0.036 | 0.042 | 0.028 | 0.040 |  |  |  |  |  |  |  |  |  |  |  |
| 7. KT879888.1_Euploea sylvester | 0.151 | 0.040 | 0.042 | 0.034 | 0.040 | 0.030 |  |  |  |  |  |  |  |  |  |  |
| 8. KR150622.1_Euploea sylvester | 0.146 | 0.038 | 0.034 | 0.036 | 0.032 | 0.028 | 0.010 |  |  |  |  |  |  |  |  |  |
| 9. H250826-025_M06_JS3_Tronga nicevillei | 0.156 | 0.034 | 0.044 | 0.002 | 0.042 | 0.030 | 0.036 | 0.038 |  |  |  |  |  |  |  |  |
| 10. H250826-025_I06_JS1_Tronga nicevillei | 0.153 | 0.032 | 0.042 | 0.000 | 0.040 | 0.028 | 0.034 | 0.036 | 0.002 |  |  |  |  |  |  |  |
| 11. ON075551.1_Euploea core | 0.153 | 0.032 | 0.042 | 0.000 | 0.040 | 0.028 | 0.034 | 0.036 | 0.002 | 0.000 |  |  |  |  |  |  |
| 12. GU012616.1_Euploea core | 0.153 | 0.032 | 0.042 | 0.000 | 0.040 | 0.028 | 0.034 | 0.036 | 0.002 | 0.000 | 0.000 |  |  |  |  |  |
| 13. LC819331.1_Euploea core | 0.153 | 0.032 | 0.042 | 0.000 | 0.040 | 0.028 | 0.034 | 0.036 | 0.002 | 0.000 | 0.000 | 0.000 |  |  |  |  |
| 14. PV592280.1_Euploea crameri nicevillei | 0.153 | 0.032 | 0.042 | 0.000 | 0.040 | 0.028 | 0.034 | 0.036 | 0.002 | 0.000 | 0.000 | 0.000 | 0.000 |  |  |  |
| 15. MH269417.1_Euploea crameri nicevillei | 0.153 | 0.032 | 0.042 | 0.000 | 0.040 | 0.028 | 0.034 | 0.036 | 0.002 | 0.000 | 0.000 | 0.000 | 0.000 | 0.000 |  |  |
| 16. PV409685.1_Euploea crameri nicevillei | 0.153 | 0.032 | 0.042 | 0.000 | 0.040 | 0.028 | 0.034 | 0.036 | 0.002 | 0.000 | 0.000 | 0.000 | 0.000 | 0.000 | 0.000 |  |

## References

Ahmed T and Mondal AC. 2016. Five butterflies new to Sundarbans Bangladesh site: how many more left to be recorded? 20th National Conference and Annual General Meeting 2016, Zoological Society of Bangladesh, Department of Zoology, University of Dhaka.

[BFD] Bangladesh Forest Department, Deutsche Gesellschaft für Internationale Zusammenarbeit (GIZ) GmbH and Jibon-Bikash Karjocrom. 2018. Butterflies of the Sundarbans: Chandpai and Sarankhola Ranges. The Dhaka Printers, Dhaka, Bangladesh.

Chowdhury S. 2014. Butterflies of Sundarban Biosphere Reserve, West Bengal, eastern India: a preliminary survey of their taxonomic diversity, ecology and their conservation. J Threat Taxa. 6(8):6082–6092. 10.11609/jott.o3787.6082-92

Chowdhury SH. 2004. Euploea crameri nicevillei (Moore, 1890) rediscovered. Bangladesh J Zool. 32(2):253–254.

Chowdhury SH 2015. Euploea crameri nicevillei. In: IUCN Bangladesh, editor. Red List of Bangladesh: Butterflies. Vol. 7. IUCN Bangladesh Country Office. p 57.

Corbet AS. 1942. Revisional notes on the genus *Euploea* F. Ann. Mag. Nat. Hist. 9:253–257.

Cramer P. 1780. De uitlandsche kapellen voorkomende in de drie waereld-deelen, Asia, Africa en America. Vol. 3. Chez S. J. Baalde. p 17–21. pl 266.

Das A, Pathania PC, Shah SK. 2019. Reporting of *Discophora sondaica* (Lepidoptera: Papilionoidea: Nymphalidae) from Sundarbans, West Bengal, India along with an updated species list from the region. Rec. Zool. Surv. India. 119(2):120–127. 10.26515/rzsi/v119/i2/2019/141450

Evans WH. 1932. The identification of Indian butterflies. 2nd ed. Bombay Natural History Society.

Fruhstorfer H. 1904. Neue Rhopaloceren aus dem malayischen Gebiet. In: Stichel, H, editor. Berliner Entomologische Zeitschrift. Vol. 49(1/2). R. Friedländer & Sohn. p 165–169. https://www.biodiversitylibrary.org/page/8345283#page/215/mode/1up

Fruhstorfer H. 1910. Familie: Danaidae. In: Seitz A, editor. Die Grossschmetterlinge der Erde: eine systematische Bearbeitung der bis jetzt bekannten Grossschmetterlinge. Vol. 9. Alfred Kernen. p 191–272.

Guindon S, Guindon S, Dufayard JF, Lefort V, Anisimova M, Hordijk W, Gascuel O. New algorithms and methods to estimate maximum-likelihood phylogenies: assessing the performance of PhyML 3.0. 2010. Systematic biology. 59(3):307–21. 10.1093/sysbio/syq010

Hall TA. 1999. BioEdit: A user-friendly biological sequence alignment editor and analysis program for Windows 95/98/NT. Nucleic Acids Sym. Ser. 41: 95–98.

Harris RB, Alström P, Ödeen A, Leaché AD. 2018. Discordance between genomic divergence and phenotypic variation in a rapidly evolving avian genus (*Motacilla*). Mol Phylogenet Evol. 120: 183–195. 10.1016/j.ympev.2017.11.020

Hebert PDN, Penton EH, Burns JM, Janzen DH, Hallwachs W. 2004. Ten species in one: DNA barcoding reveals cryptic species in the neotropical skipper butterfly *Astraptes fulgerator*. Proceedings of the National Academy of Sciences USA, 101 (41): 14812–14817. 10.1073/pnas.0406166101

Hoang DT, Chernomor O, von Haeseler A, Minh BQ, Vinh LS. 2017. UFBoot2: Improving the Ultrafast Bootstrap approximation. MBE. 35(2): 518–522. 10.1093/molbev/msx281

Hossain MM. 2013. Butterflies in the Sundarban. In: Khan R, editor, Sundarbans: Rediscovering Sundarban, the mangrove beauty of Bangladesh. Nymphaea Publication. p 100–104.

Hossain MM. 2014. Checklist of butterflies of the Sundarbans mangrove forest, Bangladesh. J. Entomol. Zool. Stud. 2(1):29–32.

Hossain MM. 2023. A review on the diversity of butterfly (Insecta: lepidoptera) fauna from Bangladesh. Bangladesh J. Zool. 51(1): 03–34. 10.3329/bjz.v51i1.68452

Hulstaert RPG. 1931. Lepidoptera Rhopalocera fam. Danaididae subfam. Danaidinae & Tellervinae. Gen. Ins.193: 105.

Inayoshi Y. 2025. Euploea crameri bremeri C. Felder & R. Felder, 1860. In: A check list of butterflies in Indo-China, chiefly from Thailand, Laos & Vietnam; [Accessed date September 30, 2025]. https://yutaka.it-n.jp/dan/30520010.html

[IUCN] International Union for Conservation of Nature. 2015. Butterflies. In: IUCN Bangladesh, editor. Red List of Bangladesh. Vol. 7. IUCN Bangladesh Country Office, Dhaka, Bangladesh.

IUCN Bangladesh. 2014. The Festschrift on the 50th Anniversary of The IUCN Red List of Threatened Species, Dhaka, Bangladesh. IUCN, x+192 pp.

Kearse M, Moir R, Wilson A, Stones-Havas S, Cheung M, Sturrock S, Buxton S, Cooper A, Markowitz S, Duran C, Thierer T. Geneious Basic: an integrated and extendable desktop software platform for the organization and analysis of sequence data. Bioinformatics. 2012 Jun 15;28(12):1647–9. 2012. Bioinformatics. 28(12): 1647–1649. 10.1093/bioinformatics/bts199

Kimura M. 1980. A simple method for estimating evolutionary rates of base substitutions through comparative studies of nucleotide sequences. J. Mol. Evol. 16(2): 111–120. 10.1007/bf01731581

Kumar S, Stecher G, Li M, Knyaz C, Tamura K. 2018. MEGA X: Molecular evolutionary genetics analysis across computing platforms. MBE. 35(6): 1547–1549. 10.1093/molbev/msy096

Lambkin TA, Braby MF, Eastwood RG, Zalucki MP. 2019. Taxonomic revision of the *Euploea alcathoe* complex (Lepidoptera: Nymphalidae) from Australia and New Guinea. Austral Entomol. 58(1): 52–75. 10.1111/aen.12299

Larsen TB. 2004.Butterflies of Bangladesh: An annotated checklist. IUCN Bangladesh Country Office. p 78. pl 5.

Mace GM. 2004. The role of taxonomy in species conservation. Philo. Trans. R. Soc. B, Biol. Sci. 359: 1444: 711–719. 10.1098/rstb.2003.1454

Moore F. 1890.Lepidoptera Indica: Rhopalocera, family Nymphalidae, sub-families Euploeinae and Satyrinae. L. Reeve & Co.

de Nicéville L. 1901.Notes on the butterflies comprised in the subgenus Tronga of the genus Euploea. JASB. 70(2): 12–38.

Rambaut A. 2018. Figtree. Version 1.4.4. Institute of Evolutionary Biology, University of Edinburgh, Edinburgh. https://tree.bio.ed.ac.uk/software/figtree/

Robinson G. 1976. The preparation of slides of Lepidoptera genitalia with special reference to the Microlepidoptera. Ent. Gaz. 27: 127–132.

Rodrigues A, Pilgrim J, Lamoreux J, Hoffmann M, Brooks T. 2005. The value of the IUCN Red List for conservation. Trends Ecol Evol. 21(2): 71–76. 10.1016/j.tree.2005.10.010

Singh N. 2025. Taxonomy in Trouble—An impediment to Life on Earth. Zootaxa. 5642(3): 395–400. 10.11646/zootaxa.5642.3.8

Talbot G. 1947. Butterflies. In: Sewell RBS, editor. The fauna of British India, including Ceylon and Burma. Vol. II. Taylor and Francis.

Thomson SA, Pyle RL, Ahyong ST, Alonso-Zarazaga M, Ammirati J, Araya JF, Ascher JS, Audisio TL, Azevedo-Santos VM, Bailly N, Baker WJ. 2018.Taxonomy based on science is necessary for global conservation. PLoS biology. 16(3):e2005075. 10.1371/journal.pbio.2005075

Xia X, Lemey P. 2009. Assessing substitution saturation with DAMBE. In: Lemey P, Salemi M, Vandamme A-M, editors. The phylogenetic handbook: A practical approach to phylogenetic analysis and hypothesis testing. Cambridge University Press. p 615–630. 10.1017/CBO9780511819049.022

Xia X, Xie Z, Salemi M, Chen L, Wang Y. 2003. An index of substitution saturation and its application. Mol Phylogenet Evol. 26(1): 1–7. 10.1016/s1055-7903(02)00326-3

